# Concurrent MEG-articulography for investigating neuromotor control of speech articulation

**DOI:** 10.1101/2020.02.05.934489

**Authors:** Blake Johnson, Qinqing Meng, Ioanna Anastasopoulou, Louise Ratko, Tunde Szalay, Michael Proctor, Cecilia Jobst, Pascal van Lieshout, Douglas Cheyne

## Abstract

Articulography and functional neuroimaging are two major tools for studying the neurobiology of speech production. Until now, however, it has generally not been possible to use both in the same experimental setup because of technical incompatibilities between the two methodologies. Here we describe results from a novel articulography system dubbed Magneto-articulography for the Assessment of Speech Kinematics (MASK), used for the first time to obtain kinematic profiles of oro-facial movements during speech together with concurrent magnetoencephalographic (MEG) measurements of neuromotor brain activity. MASK was used to characterise speech kinematics in a healthy adult, and the results were compared to measurements from the same participant with a conventional electromagnetic articulography (EMA) setup. We also characterised speech movement kinematics with MASK in a group of ten typically developing children, aged 8-12 years. Analyses targeted the gestural landmarks of the utterances /ida/, /ila/ and reiterated productions of /pataka/. These results demonstrate that the MASK technique can be used to reliably characterise movement profiles and kinematic parameters that reflect development of speech motor control, together with MEG measurements of brain responses from speech sensorimotor cortex. This new capability sets the stage for cross-disciplinary efforts to understand the developmental neurobiology of human speech production.

## 1 Introduction

Articulography and neuroimaging are the two main classes of techniques presently used by speech scientists to study the mechanisms of human speech motor control. Articulography techniques include optical tracking, ultrasound and electromagnetic articulography (EMA). The latter provides detailed measurements of movements of the peripheral speech articulators, especially the tongue, lips and jaw, and access to derived kinematic parameters including movement velocity and acceleration. In contrast, neuroimaging techniques, including functional magnetic resonance imaging (fMRI) and magnetoencephalography (MEG) provide measurements of the central brain activities associated with movements of the articulators.

Articulography and neuroimaging methods as presently available are largely incompatible with one another and cannot be used in the same experimental setup. For example, EMA uses coils that produce electrical signals when moved within a magnetic field, and both the coils and the fields’ transmitter are incompatible with the requirements of fMRI and MEG scanners. As a consequence the two types of methods must typically be used in separate experiments, and in fact have conventionally been developed and applied in quite separate academic and scientific disciplines: Articulography has been the preferred method of speech science, experimental phonology and speech language pathology; while neuroimaging is a preferred technique in neurolinguistics and cognitive neuroscience. Hence neuroimaging studies have not been able to make use of the detailed information about speech movements of the major articulators provided by articulography, relying instead on simple indices like speech onsets that provide only faint and gross indications of the precise movement trajectories of individual articulators. Conversely, articulography measurements have no access to information about the neural activities that generate and control speech movements. The neuroimaging and articulographic aspects of speech production have therefore developed to date as separate and largely independent literatures.

Recent advances in our understanding of speech motor control indicate that it would be advantageous to have access to both types of information in studies of speech production. Most notably, a study by Chartier et al. (2018) used ultrasound and video recording of speech movements in conjunction with invasive electrocorticography (ECoG) measurements of neural activity in speech motor cortex of human patients prior to surgery for intractable epilepsy. This study reported that speech motor cortex primarily encodes information about kinematic parameters derived from the speech tracking measurements, rather than acoustic or phonemic parameters derivable from the acoustic speech signal. Such findings conform well to concepts within state speech motor control models such as Articulatory Phonology (AP) and the associated Task Dynamics framework, which hold that articulators create functional relationships in order to cause local vocal tract constrictions (Goldstein & Fowler, 2003). The abstract representations of these articulatory events during speech production are called gestures, the basic units of phonological contrasts (Browman & Goldstein, 1992). Gestures are individual and context-invariant units which can be combined into larger sequences such as syllables, words and phrases to create meaningful language-specific contrasts. Moreover, gestures are task-specific vocal tract actions which can be implemented by coordinated activity of the articulators in a contextually appropriate manner (Van Lieshout, Merrick, & Goldstein, 2008). According to Gafos (2002), gestures are described as dynamic spatio-temporal units. In other words, a gesture can be described as “a member of a family of functionally equivalent articulatory movement patterns that are actively controlled with reference to a given speech-relevant goal” (Saltzman & Munhall, 1989).

There are thus good theoretical and empirical reasons to believe that both articulographic and neuroimaging information are needed to advance our understanding of speech motor control. MASK was designed to track speech and nonspeech movements during magnetoencephalographic measurements of brain function (Alves et al., 2016). The MASK system is integrated with the MEG and it consists of three components: tracking coils, coil driver electronics, and a coil-tracking algorithm. In contrast to the passive induction coils used in EMA, MASK coils are actively powered non-magnetic inductor chips that emit high-frequency oscillating magnetic fields which are detected by the MEG sensors and spatially localised. These signals are then separated from the MEG signals by low-pass filtering, providing continuous motion signals time-locked to the brain activity data without interfering with the brain measurements. The MASK system can track movement at rates of up to 50 cm/s with less than 1 mm relative position error (Alves et al., 2016). Importantly, this system does not require line-of-sight tracking, allowing for measurements from all oral articulators including the tongue.

### 1.1 Aims

In the present study we aimed to (1) compare MASK measurements of speech kinematics in a healthy adult to measurements on the same individual with a conventional electromagnetic articulography (EMA); and (2) Characterise speech movement kinematics with MASK in a group of ten typically developing children. Two main types of non-linguistic speech utterance protocols were employed. In the first, participants produced /ida/ and /ila/ in a self-paced fashion at a rate of about one utterance every four seconds. Analyses targeted the gestural landmarks of the consonants in these two utterances. In the second, participants produced /pataka/ in a speeded fashion and analyses examined the consistency/variability of tongue dorsum, body and tip movements over repeated utterances. While the scope of the present report is limited to the articulographic aspects of the MASK technology, we consider implications for neuroimaging studies of speech motor control in the discussion section.

## 2 Methods

### 1.1. Participants

Participants were recruited from the Sydney area and all were native speakers of Australian English. All procedures protocols were approved by the Macquarie University Human Research Ethics Committee. The adult participant was a healthy male aged 48 years with no history of speech, language or hearing difficulties.

Inclusion criteria for children were a) age 8 – 12 years; b) no speech, language or hearing difficulties; c) native Australian monolingual English speakers; d) no orthodontic or metal implants; e) no history of head injury; f) no medications that might affect motor or cognitive performance; g) no known neurological, social-emotional (affective) deficits, h) no structural vocal tract issues; i) no visual problems; j) no speech prosody and voice issues.

Children visited the lab twice accompanied with their parents/caregivers. During their first visit, parents completed a developmental and medical case history form to ensure that none of the participants have known speech, language, neurological and cognitive deficits. MEG brain measurements and MASK speech tracking measurements were carried out in a second visit.

Children were screened with speech, language and hearing assessments by a certified speech-language pathologist. All screened children performed within age appropriate limits in speech, expressive and receptive language and oral-motor assessments including the Clinical Evaluation of Language Fundamentals-Fifth Edition, Screening Test Australian & New Zealand Language Adapted Edition (CELF -5 Screening Test; Wiig et al., 2013), the Oral Speech Mechanism Screening Evaluation-Third Edition (St. Louis & Ruscello, 2000) and the Goldman-Fristoe Test of Articulation 2 (GFTA-2) (Goldman R, 2000). All children had normal pure tone hearing thresholds. Handedness was assessed using the Edinburgh Handedness Inventory-Short Form (Veale, 2014).

### 2.1 Experimental Tasks

During the experiment, participants were instructed to fix their gaze on a central cross projected to a ceiling screen while producing two blocks of simple speech tokens “/ida/” and “/ila/” in a self-paced fashion, with a duration of 240 seconds for each block and approximately 2 seconds inter-production interval. Detailed speech related movement from selected articulator locations were acquired with MASK and recorded concurrently with the speech audio signal and MEG brain activity.

### 2.2 Experimental protocol

Data collection was preceded by a training and familiarization session in a mock MEG scanner (Rapaport et al., 2019). Participants were trained to ensure familiarity with the procedures, to avoid incorrect speech productions or head movements, and to familiarise the children with the environment, the experimental setup and the researchers. Each participant was required to produce each task correctly before the data acquisition began. During training, the experimenter decided whether children’s speech productions were accurate and whether the procedure had been fully understood. If not, the stimuli were repeated and further instructions were given until participants produced the utterances correctly and at the correct rate.

After training, five head position indicator (HPI) coils were secured to the head by an elasticized head cap. Participant’s head shapes and fiducial positions were digitised (Polhemus Fastback, Colchester, VT).

All participants performed three speech production tasks (summarised in **Table 1**). Productions included the monosyllabic nonwords /ida/ and /ila/ produced in a self-paced rate. The vowel-consonant-vowel (VCV) nonwords consisted of either the voiced, alveolar, stop consonant /d/, or the voiced, alveolar, lateral approximant /l/ combined with high, front and low, back vowels /i/ and /a/ respectively. The voiced, alveolar stop /d/ is produced by the transient occlusion of the vocal tract by the tongue tip whereas the lateral consonant /l/ is produced by the anterior constriction of the tongue tip and the posterior constriction of the tongue dorsum, with the lateral sides of the tongue lowered. These consonants were selected because they differ in terms of their articulatory complexity (/d/ is an early acquired sound and articulatorily relatively simple, while /l/ is a late acquired sound and articulatorily relatively complex) but with the same central place of articulation (both alveolar) allowing us to compare their kinematic properties. The vowel /i/ is produced by a high, front tongue constriction during phonation and the vowel /a/ is produced by a low, back tongue retraction. Some consonant segments consist of a single gesture, while some others are more complex and require the coordination of more than one gestures (Studdert-Kennedy & Goldstein, 2003). Concurrently, sounds which are acquired early in speech development, require the coordination of independent gestures (such as /d/ involving tongue tip and jaw), while late-acquired sounds (such as /l/) require the coordination of two lingual gestures (Gick et al., 2007). In particular, /d/ is produced by the transient occlusion of the vocal tract by the tongue tip while /l/ is characterised by an anterior constriction of the tongue tip in the alveolar ridge which allows for lateral airflow (Gick, Wilson, & Derrick, 2012) and by a posterior constriction of the tongue dorsum which is either pharyngeal or velar (Lin & Demuth, 2015).

**Table 1.** Description of targeted productions.

| Nonword | Vowel | Consonant |
| --- | --- | --- |
| ida | high front<br>low back | voiced, alveolar, stop |
| ila | high front<br>low back | voiced, alveolar, lateral approximant |
| pataka | low back | voiceless, bilabial, stop<br>voiceless, alveolar, stop<br>voiceless, velar, stop |

**Table 2.** Mean (standard deviation) values of gestural movements for /ida/ and /ila/. Calculated for the tongue tip coil for /d,l/ consonant productions.

|  | Consonant /d/ |  |  | Consonant /l/ |  |  |
| --- | --- | --- | --- | --- | --- | --- |
| DISTANCE [in mm] |  |  |  |  |  |  |
|  | CL | TAR | OP | CL | TAR | OP |
| EMA | 6.42 | 0.46 | 11.88 | 3.71 | 0.64 | 7.32 |
| Adult | (1.34) | (0.46) | (2.38) | (0.47) | (0.46) | (1.05) |
| MASK | 3.29 | 1.32 | 8.19 | 2.21 | 0.6 | 5.3 |
| Adult | (0.95) | (1.05) | (1.86) | (0.65) | (0.48) | (1.01) |
| MASK | 4.05 | 1.54 | 8.64 | 2.36 | 1.06 | 8.04 |
| Children | (2.89) | (1.12) | (3.96) | (1.64) | (0.69) | (3.43) |
| DURATION [in msec] |  |  |  |  |  |  |
|  | CL | TAR | OP | CL | TR | OP |
| EMA | 96.58 | 59.34 | 107.32 | 84.63 | 58.07 | 109.48 |
| Adult | (18.8) | (8.93) | (8.7) | (7.61) | (8.5) | (7.3) |
| MASK | 144.3 | 111.85 | 180.82 | 126.6 | 107.31 | 170.12 |
| Adult | (17.5) | (25.5) | (16) | (17.6) | (18) | (18.9) |
| MASK | 206.5 | 102.46 | 229.15 | 164.55 | 117,33 | 237,8 |
| Children | (50.85) | (32,75) | (50.54) | (44.83) | (31,63) | (49,06) |
| PEAK VELOCITY [in mm/s] |  |  |  |  |  |  |
|  | CL |  | OP | CL |  | OP |
| EMA | 10.38 |  | 18.52 | 8.13 |  | 14.99 |
| Adult | (1.48) |  | (3.07) | (1.44) |  | (1.39) |
| MASK | 3.31 |  | 6.02 | 2.54 |  | 5.05 |
| Adult | (0.84) |  | (1.51) | (0.71) |  | (0.94) |
| MASK | 3.44 |  | 6.35 | 2.45 |  | 5.33 |
| Children | (1.25) |  | (1.63) | (0.82) |  | (1.63) |

CL: closing movement, TAR: target plateau, OP: opening movement.

A third task involved repetitive productions of the tri-syllabic nonword [pataka]. This stimulus was selected for measuring intra- (between single articulator movements) and inter (between consonant and vowel gestures) gestural coordination within a single task (van Lieshout et al., 2007). The approach used in this investigation is similar to that used by other researchers as in the study of Van Lieshout & Moussa (2000) where participants produced them in single trials after taking a deep breath. In the same line, Van Lieshout et al. (1999) investigated the coordination dynamics of the coupling between upper and lower lip in bilabial gestures in four adult participants. The experimental tasks consisted of reiterant speech tasks and participants were producing them in single trials of 10s repeating as many times as they could. Nip, (2015) focus on the interarticulator coordination of children with cerebral palsy and controls by utilising maximum performances tasks which differed in terms of their linguistic and articulatory demand. In this study, participants had to produce the DDK task as quickly as they could in a single breath remaining intelligible. Van Lieshout et al., (2007) used disyllabic and trisyllabic reiterant nonwords in a single trial and at preferable rate in order to study motor coordination in an adult with apraxia of speech (AOS) and a group of age-matched controls.

### 2.3 Data acquisition

#### 2.3.1 EMA

Articulatory data of the adult participant were collected using electromagnetic articulography (NDIWave, Waterloo, Canada) and were recorded at a sampling rate of 100 Hz. Sensor coils were attached in a mid-sagittal placement to the lower incisor, upper and lower lip, tongue tip, tongue body and tongue dorsum. Three additional coils were attached to the bridge of the nose (nasion) and to the the left and the right mastoids for head motion correction.

After the coils were attached, the occlusal bite plane was measured with a plastic protractor with three coils attached. The participant held the device in their mouth using their teeth for about 10 seconds. The midline of the participant’s palate was traced with a custom 6D palate probe (NDI, Waterloo, Canada).

The participant was seated in a straight back chair and was recorded while reading the nonword stimuli which were presented on a computer screen. Ten productions of the same nonwords which were used in the MASK experiment were produced following the same experimental protocol. Audio recordings were obtained using a microphone (Røde Model NT1-A, Long Beach, USA) placed 40 cm from the participant’s lips and digitized with a 22050 Hz sampling rate.

#### 2.3.2 MASK

During the MEG portion of the experiment, MASK tracking coils were attached to the midline positions of the vermilion border of upper and lower lip, the lower incisors, the tongue tip, the tongue body and the tongue dorsum. MASK tracking coils are attached with their magnetic moment oriented either parallel or perpendicular to the surface of different articulators (e.g. tongue versus lip) such that the magnetic field lines are always directed towards the MEG sensors. The tongue sensors were attached to the tongue of the children with EPIGLU® surgical glue. The lower incisor sensor was attached to a thin thermoplastic mould used to cover the lower incisors of the participants (Van Lieshout & Moussa, 2000). Surgical tape was used to affix the upper and lower lip sensors as well as the three coils attached to the nasion, the left and the right preauricular points. Tongue coils were affixed to the tongue tip/blade (1 cm from the tongue tip), tongue body (2 cm from the tongue tip) and tongue dorsum (4 cm from the tongue tip) (**Figure 1**). The occlusal plane and head alignment were measured using a plastic protractor. Participants were instructed to place the protractor between the upper and lower teeth and bite it during the digitisation process. The protractor was placed such that the midline was aligned with the mid-sagittal plane. The nose bridge, the left, the right preauricular and three points (triangular formation) of the protractor were digitised with a pen digitizer (Polhemus Fasttrack, Colchester, VT).

**Figure 1.**
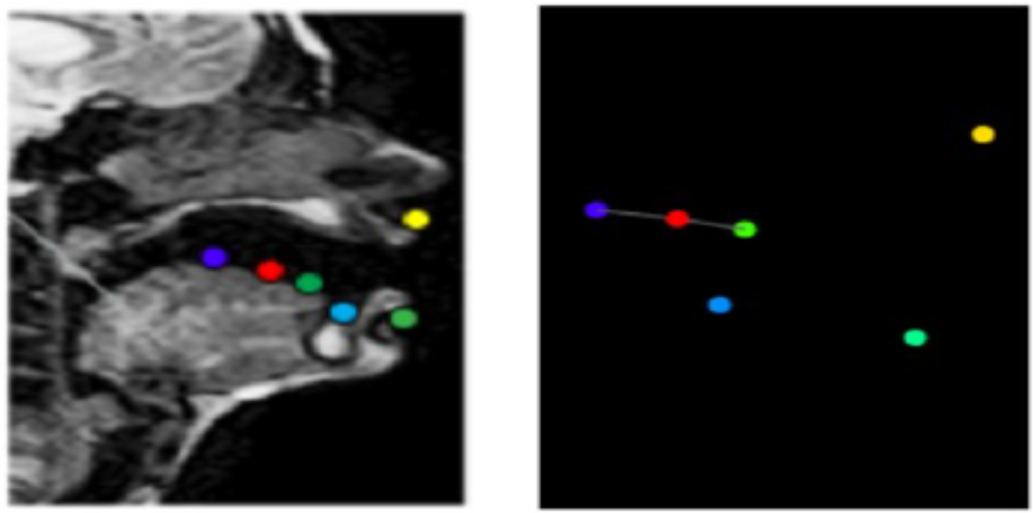
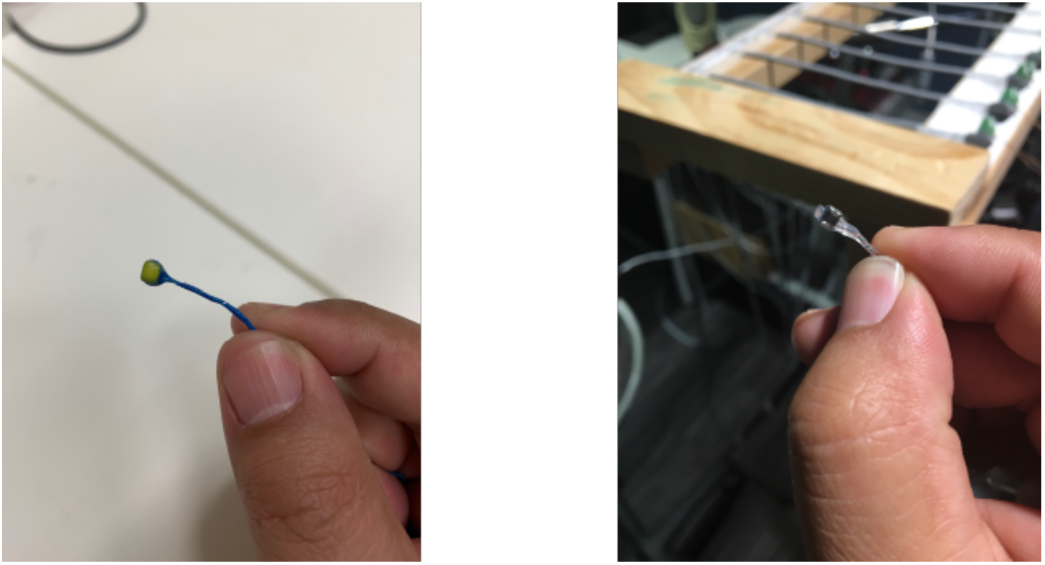
MASK coils and positioning. Top: Midsagittal MRI shows positioning of three tongue coils, lower incisor, and upper and lower lip coils. Right panel shows coil positions during speech production in MEG scanner. Bottom: MASK coil (left) and EMA coil (right).

#### 2.3.3 MEG

Brain activity and MASK tracking data were recorded continuously using a whole-head MEG system (Model PQ1160R-N2, KIT, Kanazawa, Japan), consisting of 160 first-order axial gradiometers with a 50-mm baseline (Kado et al., 1999; Uehara et al., 2003). MEG data were acquired with a bandpass of 0.03 Hz - 1000 Hz, 4000 Hz sampling rate and 16-bit quantization precision. The measurements were carried out with participants in a supine position in a magnetically shielded room (Fujihara Co. Ltd., Tokyo, Japan). Audio speech recordings were obtained with an optical microphone (Optoacoustics, Or-Yehuda, Israel) fixed at a distance of 20 cm from the mouth of the speaker and digitised using a sound card with 48 kHz sample rate and 8-bit quantization precision (Creative Labs Model X-Fi Titanium HD, Singapore). Marker coil positions were measured before and after each recording block to quantify head movements, with a maximum displacement criterion of < 5 mm in any direction. The total duration of the experiment was about 45 minutes.

Magnetic resonance images (MRI) of the head and brain were acquired using a 3 Tesla Siemens Magnetom Verio scanner with a 12-channel head coil. Images were acquired using an MP-RAGE sequence (192 axial slices, TR = 2000 ms, TE = 3.94 s, FOV = 240 mm, voxel size= 0.9 mm3, TI = 900, flip angle = 9°).

For the self paced productions of /ida/ and /ila/ participants were required to fixate on a small white cross on back-projection screen. When the cross disappeared, a text indicating the required speech production was shown on the screen to cue the beginning of the trial. Participants were instructed to produce the utterance at a self-paced rate of about one item every two seconds as long as the cue was remaining on the screen. Short breaks were provided after each trial. Participants were instructed to close their mouth after each utterance to reduce variability across repetitions. Participants completed 80-120 productions of each speech token and short breaks were provided after each condition.

For the reiterated diadochokinetic task /pataka/, participants were shown text which asked to take a deep breath and to repeat ‘as many productions as they can’ when a white cross appeared on the screen. The disappearance of the cross with a new visible cue signalled the beginning of the trial. Participants produced the nonword repetitively until the cue disappeared. Each trial lasted 10 seconds. Participants completed eight trials.

### 2.4 Analysis

#### Kinematic analyses

For MASK and EMA data the locations of reference points marked on the biteplane were used to transform all movement data in the coordinate system defined by the nasion, right and left preauricular point to a new coordinate system defined by the biteplane reference points. The tongue tip gestures of /d/ and /l/ were analysed in twenty utterances per participant (ten /ida/ and ten /ila/ tokens).

After the identification of the temporal landmarks and their values, for each consonantal gesture, three different movements were identified: *Closing movement*, when the TT moves towards its target (TT constriction for the consonants /d/ and /l/); *target plateau*: the interval of time when the closure/constriction is actively held; and *opening movement* or release when the articulator moves away from the palate (Gafos, 2002, Goozee et al., 2000). The kinematic parameters included in the current analysis were: *distance* (mm) travelled by the principal receiver coil during closing, opening movements and during the target plateau of the two consonants; *duration* (ms) of the closing, opening movement and target plateau and *peak velocity* during closing and opening movements.

MVIEW software (Gafos, Kirov & Shaw, 2005) was used for visualization and analysis of the articulatory data sampled at 100 Hz (EMA) and 25 Hz (MASK) respectively, with a discrete cosine transform (DCT) based automated smoothing procedure applied (Garcia, 2010). Gestural landmarks were labelled manually. Ten temporal landmarks were extracted for each gesture:

i. gestural onset (GONS) (ms)- the time point of the gestural onset
ii. onset peak velocity (PVEL) (ms) - the time point of maximum TT velocity during the closing movement, before the target plateau
iii. onset of target plateau (NONS) (ms)- the time point of the onset of the target plateau
iv. offset of target plateau (NOFFS) (ms)- the time point of the offset of the target plateau
v. offset peak velocity (PVEL2) (ms)- the time point of maximum TT velocity during the opening movement, after the plateau
vi. gestural offset (GOFFS) (ms) - the time point of the gestural offset.

The spatial values of the temporal landmarks were also identified: gestural onset (GONS) (mm), onset of target plateau (NONS) (mm), offset of target plateau (NOFFS) (mm) and gestural offset (GOFFS) (mm). Moreover, the speed was extracted in cm/sec for the onset PVEL and PVEL2. **Figure 2** illustrates the measurements of the kinematic parameters used.

**Figure 2.**
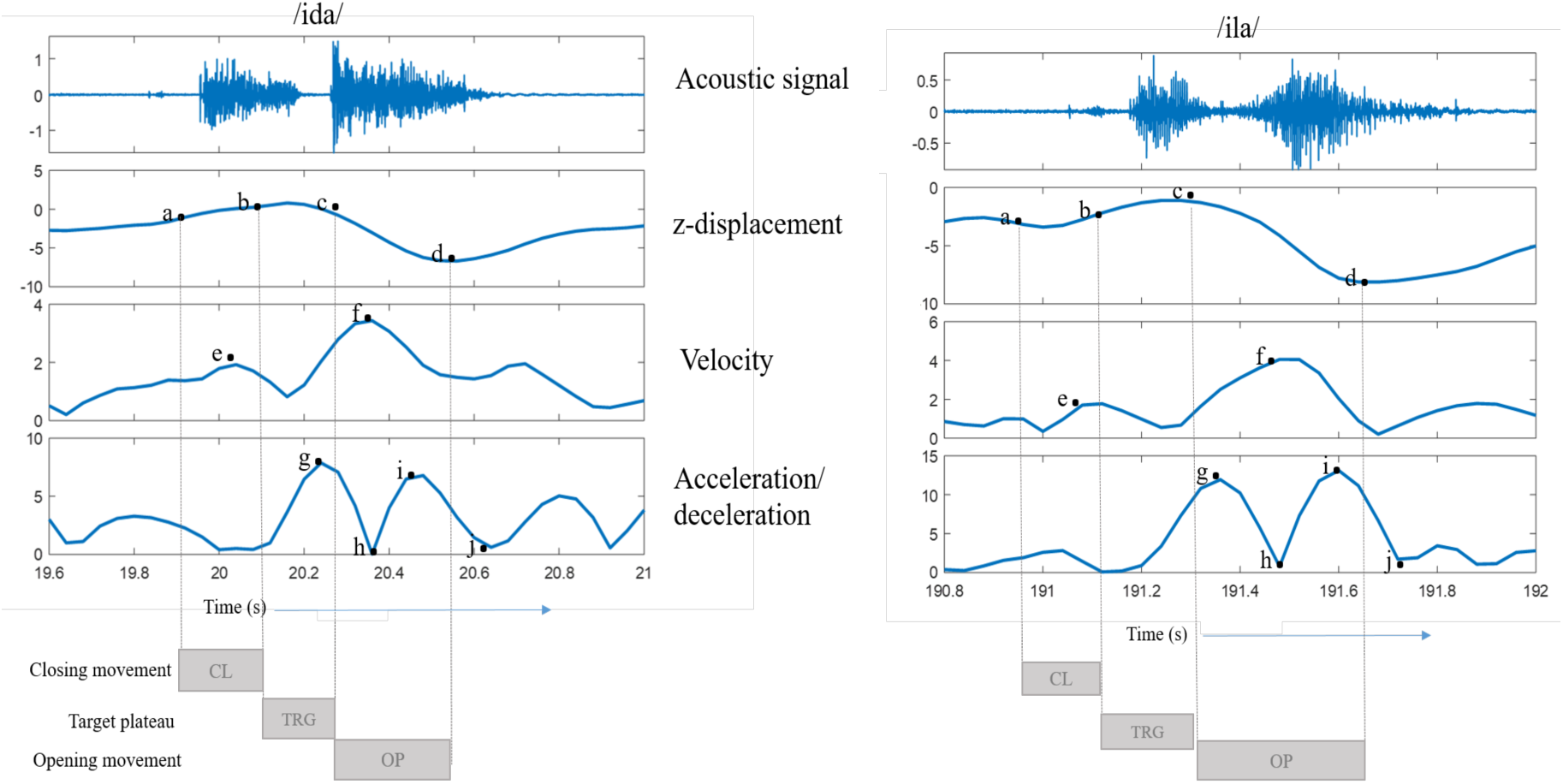
Analysis of gestures. Tongue tip displacement is shown. Vertical lines represent key articulatory landmarks: gestural onset (a), onset of target plateau (b), offset of target plateau (c) and gestural offset (d).

Variability of the tongue tip, body and dorsum motions was investigated over repetitions of the diadochokinetic rate (DDK) /pataka/. In order to segment the /pataka/ for analysis, z-displacements and velocity signals of the principal articulators were used to extract the first and the last movement of the utterance. The peak velocity of the first opening movement (release of /p/ to /a/ in /pataka/) to the peak velocity of the last opening movement (release of /k/ to /a/ in /pataka/) were selected by visual inspection of three productions of every block of repetitions with the first production of every block removed. After the start and end points were selected, the spatiotemporal index (STI; Smith et al., 1995; Smith et al., 2000) was calculated for 10 blocks of three reiterated /pataka/’s for each participant. The displacement signals were amplitude- and time-normalised and variances were computed at 2% intervals across repetitions of the normalised displacement waveforms. The resulting STI was the square root of the sum of these 50 variances (In contrast to Smith et al. (1995), we elected to sum variances -- which are additive -- rather than standard deviations).

#### 2.4.1 MEG Analysis

MEG data analysis was performed with the opensource toolboxes Fieldtrip (Oostenveld, Fries, Maris, & Schoffelen, 2011) and Brainwave (Jobst et al., 2018) and custom MATLAB scripts. Offline MEG data were first filtered with a high-pass filter (0.1 Hz), a low-pass filters (30 Hz) and a notch filter (50 Hz, 100 Hz, 150 Hz) and then segmented into epochs of 1.5 seconds in length according to the onset of speech audio recording. All data trials were down-sampled to 200 Hz prior to independent component analysis (ICA)(Makeig, Bell, Jung, & Sejnowski, 1996) to remove eye-blinks, eye-movements, heartbeat-related artefacts and magnetic jumps. Components corresponding to those artefacts were identified by their spectral, topographical and time course characteristics.

## 3 Results

Examples of raw tracking results for EMA and MASK recordings for two adult participants are shown in **Figure 3**. In general, the raw MASK tracking waveforms were visually highly similar to those obtained with EMA.

**Figure 3.**
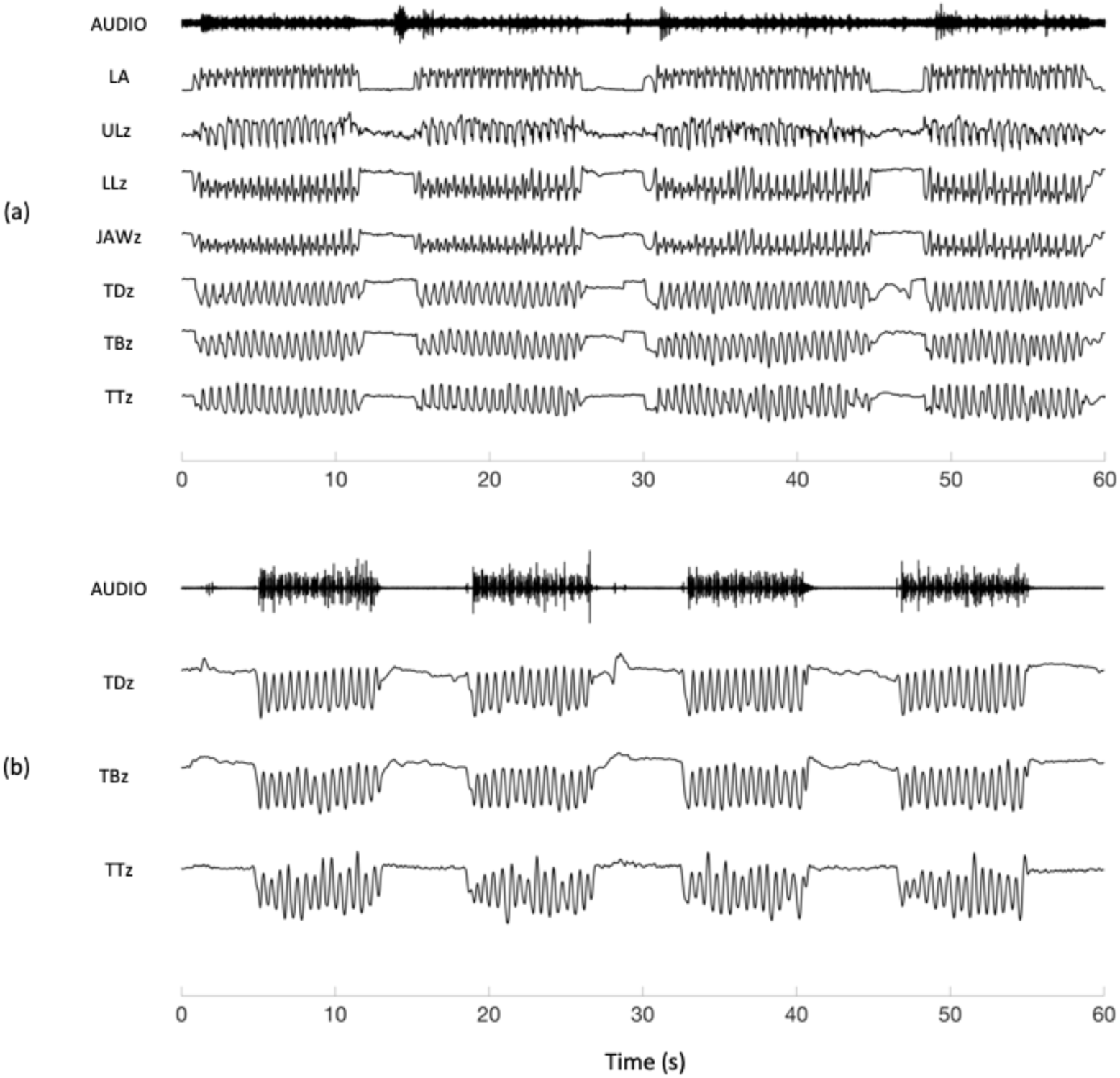
Tracking with EMA and MASK. Examples of raw tracking results in sagittal (z) direction, reiterated /pataka/ (a) EMA tracking results for the adult participant. Acoustic recording shows high noise baseline due to suboptimal placement. (b) MASK tracking result for the same adult participant for reiterated /pataka/. LA lip aperture. UL Upper Lip. LL Lower Lip. TD Tongue dorsum. TB Tongue Body. TT Tongue Tip.

All six tracking coils provided stable tracking results for the EMA recordings. For the adult participant who participated in both EMA and MASK recordings, the three MASK tongue coils provided stable tracking results throughout the recording session, while the lip and jaw coils did not. The EMA session voice recording of this participant however had a high noise baseline due to a suboptimal microphone placement (**Figure 3**, top trace).

### 3.1 Consonant gestural landmarks for the adult participant

Results described below are for consonant gestural analyses for the tongue tip sensor for 10 repetitions of /ida/ and /ila/, from the single adult who participated in both EMA and MASK recording sessions (left two panels of **Figure 4**).

**Figure 4.**
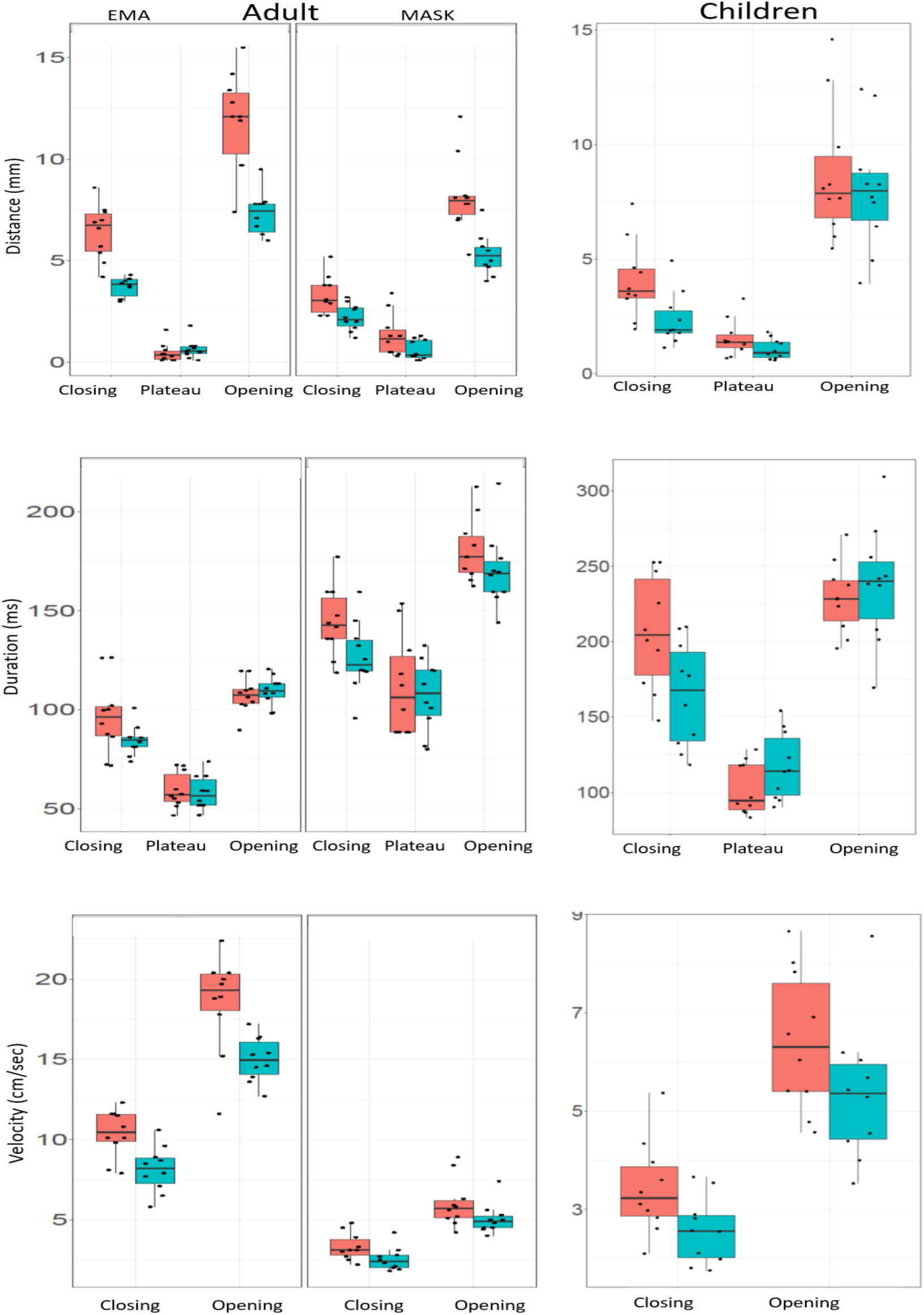
Kinematic parameters derived from EMA and MASK. Kinematic parameters from EMA and MASK. Distance, duration and peak velocity values of tongue tip of /d,l/ consonant productions of an adult participant in EMA and MASK (left) and for the group of 10 children in MASK (right). /d/ = red, /l/ = cyan.

MASK *distance* values were significantly smaller than EMA distance values for opening and closing movements for both /ida/ and /ila/ productions (2 tailed paired sample t-test, p < .001 for all). MASK *duration* values were significantly greater than EMA duration values for all movements and both consonant productions (p < 0.001 for all). Given the shorter distances and longer durations, the derived MASK *velocities* were significantly lower than those from EMA (p < 0.001 for all movements and both consonants).

Despite the lower distance and velocity, and larger duration values in MASK, the overall profiles of both consonantal gestures show the same pattern in both MASK and EMA:

- Distance and duration: closing movement > opening movement > target plateau (2 tailed, paired sample t-tests, all contrasts, p < .001)
- Velocity: closing movement > opening movement (p < .001).

Within each tracking method, significant differences (2 tailed, paired sample t-tests, Bonferroni correction = .05/16) between the /d/ and /l/ consonantal gestures were obtained for: MASK distance, closing movement (p < .001); EMA distance, opening movement (p < .001); MASK distance, opening movement (p < .001); and MASK peak velocity, closing movement (p < .001).

### 3.2 Consonant gestural landmarks for the group of children

Results described below are for consonant gestural analyses for the tongue tip sensor, for the group of 10 children. For each child, data points are means for 10 repetitions of /ida/ and /ila/.

**Figure 4** shows that the overall profiles of both consonantal gestures show the same pattern obtained for the adult participant in EMA and MASK:

- Distance and duration: closing movement > opening movement > target plateau (2 tailed, paired sample t-tests, all contrasts, p < .001)
- Velocity: closing movement > opening movement (p < .001).

None of the /d/ versus /l/ contrasts for the group of children reached statistical significance after correction for multiple comparisons (2 tailed, paired sample t-tests, Bonferroni correction = .05/8).

### 3.3 Consistency/variability of tongue movements in reiterated /pataka/ productions

The variability of the entire articulatory movement trajectory for the group of children and the adult participant was computed with the spatiotemporal index (STI). **Figure 5** illustrates the normalised displacements of 10 tokens of three successive reiterated productions of /pataka/, for the tongue dorsum, tongue body and the tongue tip for one child and one adult participant. The overlaid plots and associated STI values show that (1) most variation is seen in the tongue tip for both of these participants; and that (2) the child participant showed more variability in all three articulators.

**Figure 5.**
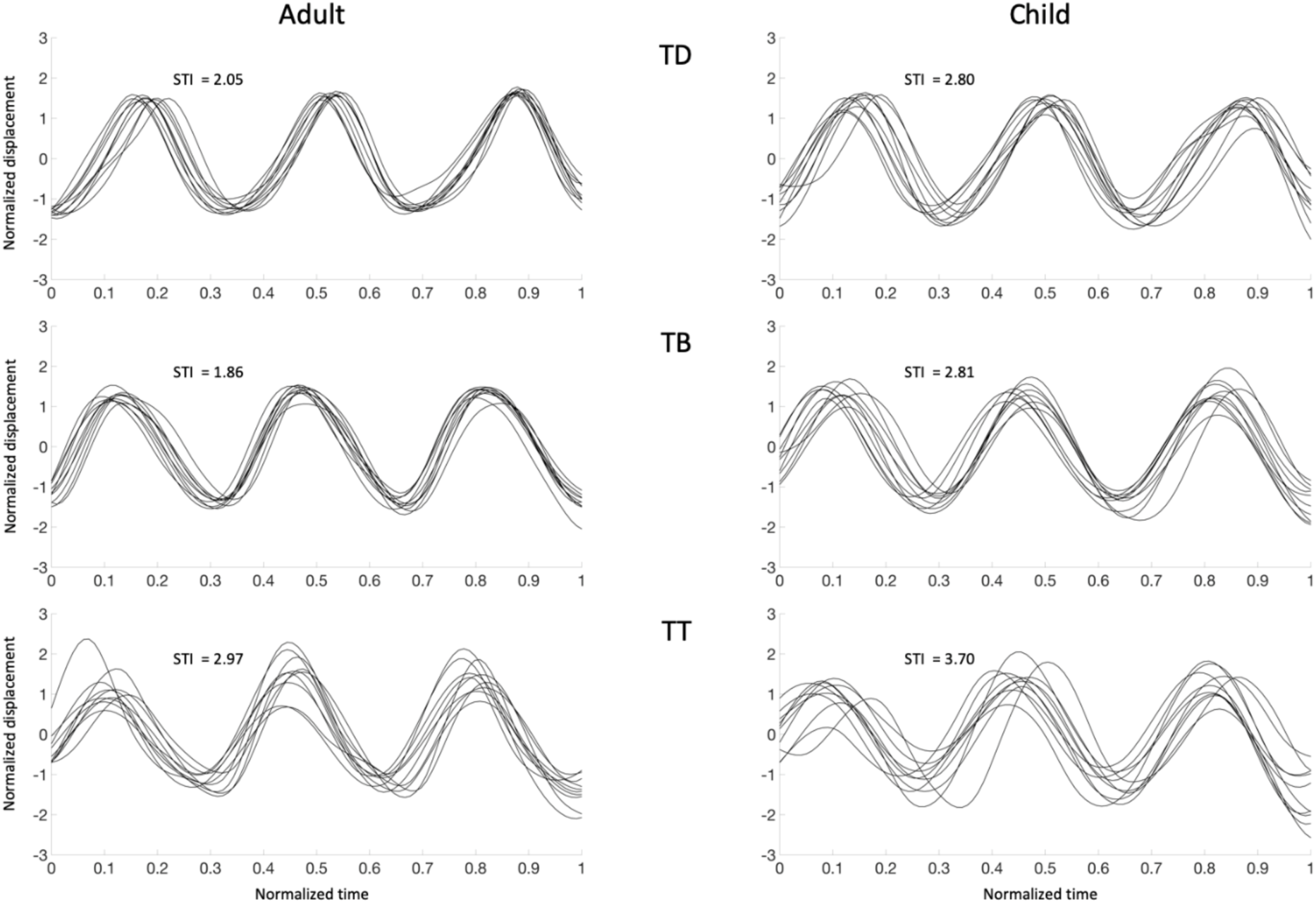
Amplitude and time normalised MASK tongue displacement plots for /pataka/ (z direction). Overlays of 10 productions of three tokens of /pataka/ for an adult and a child participant. TD = tongue dorsum; TB = tongue body; TT = tongue tip. STI = spatiotemporal index.

**Figure 6** and **Table 3** summarize STI values for the group of children relative to the STI values of the adult. The STI of tongue dorsum was calculated in 9 children as the tongue dorsum coil dropped off in one of the participants during the experiment (**Table 3**). All STI values for the adult were at the very bottom of the STI range for the group of children (note that the lowest STI values for all tongue positions were obtained from the same child, P2). Within the group of children, STI values showed small, non-significant correlations with age (TD: r = -.14; TB: r = -.14; TT: r = -.38).

**Table 3.**
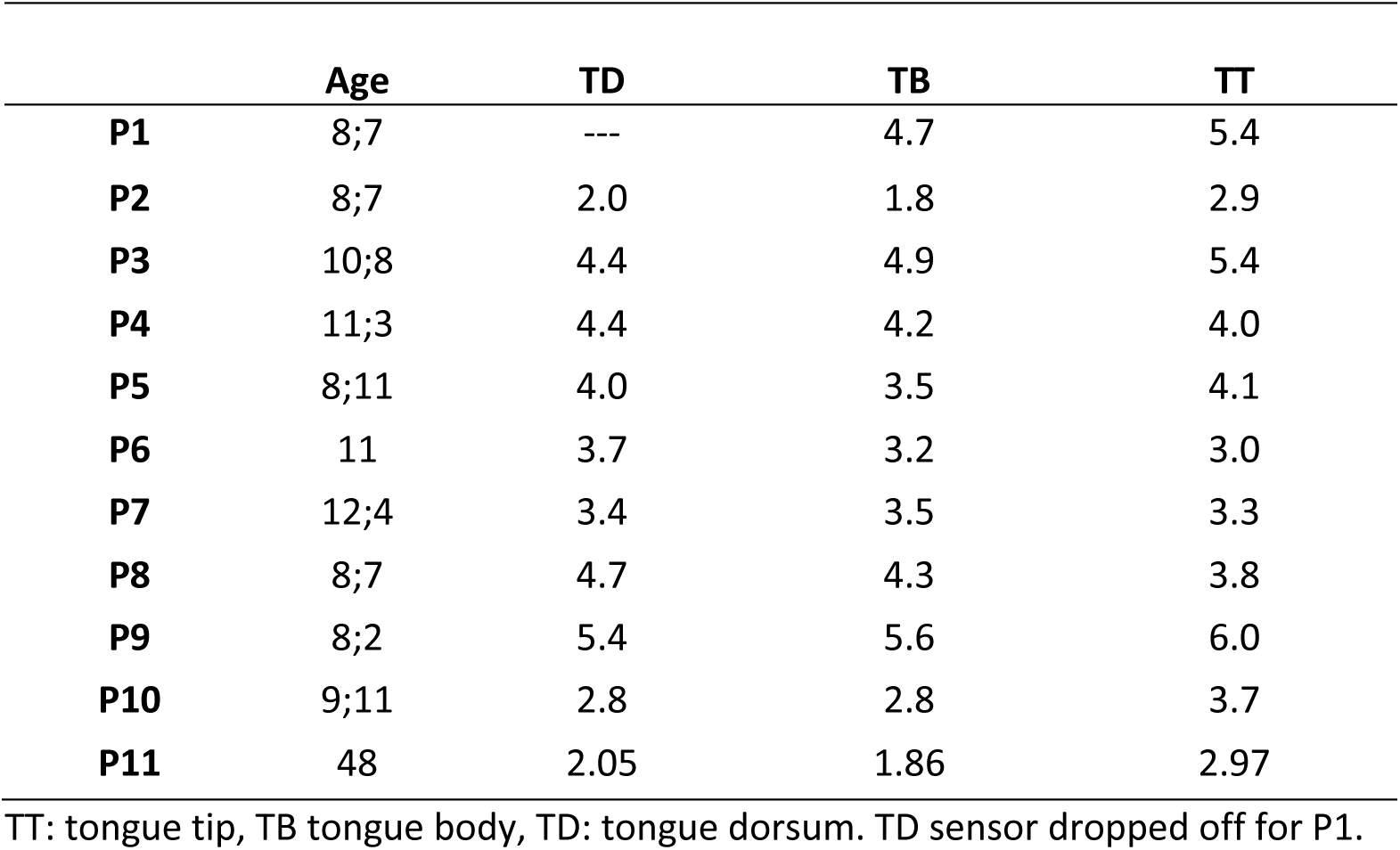
STI values for /pataka/ productions.

|  | Age | TD | TB | TT |
| --- | --- | --- | --- | --- |
| **P1** | 8;7 | --- | 4.7 | 5.4 |
| **P2** | 8;7 | 2.0 | 1.8 | 2.9 |
| **P3** | 10;8 | 4.4 | 4.9 | 5.4 |
| **P4** | 11;3 | 4.4 | 4.2 | 4.0 |
| **P5** | 8;11 | 4.0 | 3.5 | 4.1 |
| **P6** | 11 | 3.7 | 3.2 | 3.0 |
| **P7** | 12;4 | 3.4 | 3.5 | 3.3 |
| **P8** | 8;7 | 4.7 | 4.3 | 3.8 |
| **P9** | 8;2 | 5.4 | 5.6 | 6.0 |
| **P10** | 9;11 | 2.8 | 2.8 | 3.7 |
| **P11** | 48 | 2.05 | 1.86 | 2.97 |

TT: tongue tip, TB tongue body, TD: tongue dorsum. TD sensor dropped off for P1.

**Figure 6.**
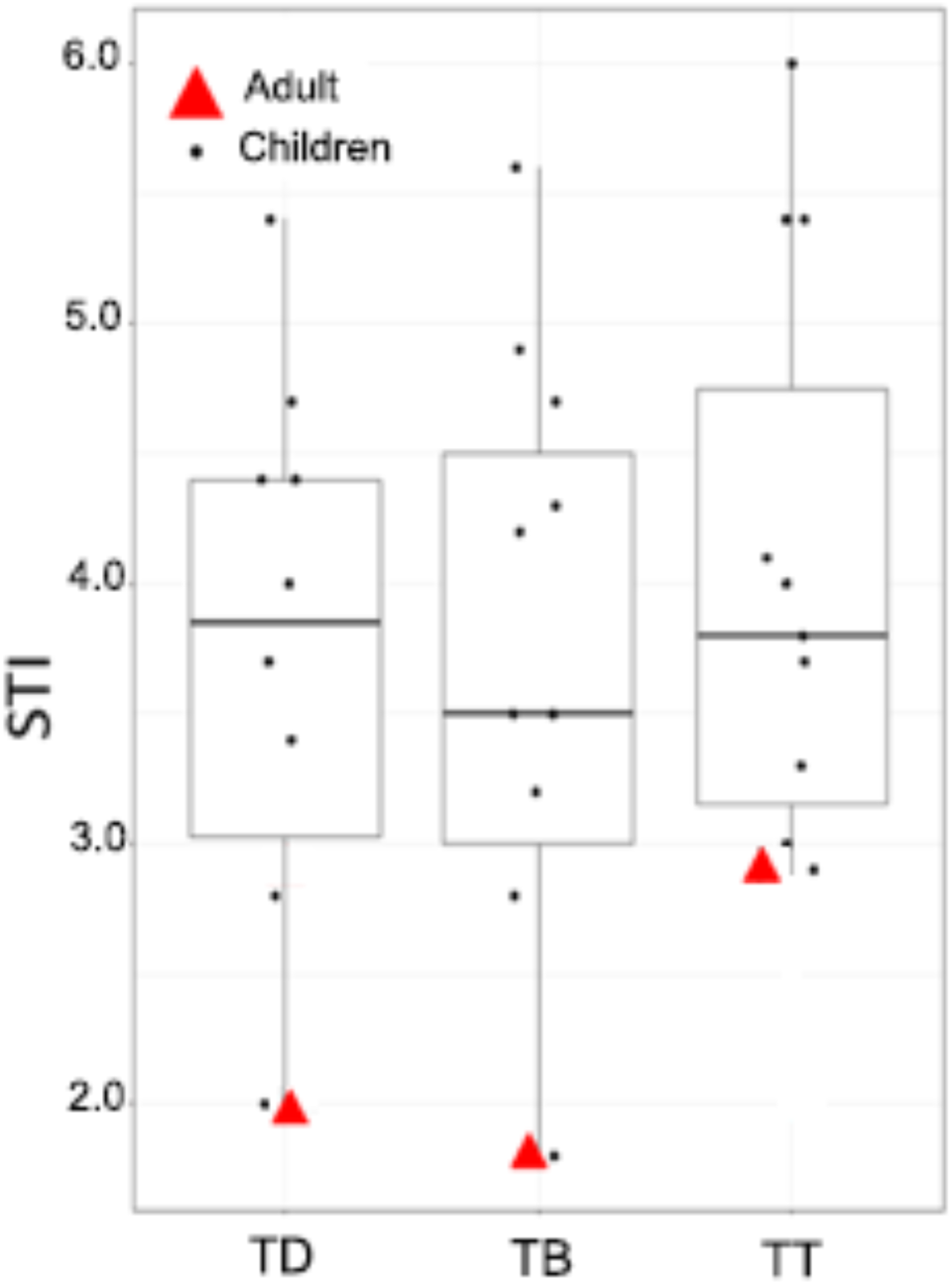
Spatiotemporal index for /pataka/ productions.

## 4 Discussion

The present report describes the initial experiences of our group using a novel articulographic approach for tracking speech movements and characterising the salient kinematic parameters of those movements within the MEG brain scanner. Detailed descriptions of an older version of the MASK system components and spatial and temporal calibrations and accuracy are provided in Lau (2013) and Alves et al. (2016).

Gestural durations derived from the present MASK setup were significantly longer, distances were smaller, and velocities were correspondingly lower than those obtained from EMA in the same participant. There are three possible sources for these discrepancies. First, the two experimental setups required quite different body postures. In EMA, the participant was seated in a straight-backed chair and facing a computer screen; in contrast, and as is common in many functional neuroimaging setups, the MEG-MASK setup required supine positioning with the participant looking up at a projector screen. Breathing control may be expected to be different in the two positions, with less ability to generate higher subglottal pressures and the tongue may be more affected by gravity toward the posterior position in the supine position. There currently exists little literature to bear on this question. Vorperian et al. (2015) reported significantly smaller vocal tract volumes in the supine position compared to upright positioning as well as significantly larger values for acoustic measurements of F0 and some formant frequencies. On the other hand, Dietsch & Cirstea (2013) reported no difference in tongue pressure across upright and supine positions. Further postural comparison studies are required to clarify this issue.

Visual examination of the tracking data indicated that MASK tracking amplitudes were typically lower in comparison to EMA tracking amplitudes, with MASK having a generally lower dynamic range for movement tracking. This would account for longer gestural durations and smaller MASK distances since MASK peaks and valleys were less sharp and well-defined. One contributing reason it was not possible to check and assess the adequacy of attachment of tracking sensors until after completion of the experiment with the current MASK setup, and this may have resulted in suboptimal placement of some sensors. As noted below, we have subsequently incorporated a capability for real-time visualization of MASK sensor tracking comparable to that used in typical EMA setups. This will substantially improve the optimization of sensor placements during the experiment.

In spite of these differences in absolute values, the overall gestural *profiles* obtained from MASK and EMA (e.g. opening movement relative to closing movement relative to plateau) were highly similar for all three kinematic parameters analysed. Further, the *relative profile differences* between /d/ versus /l/ were nearly identical for MASK and EMA, although these were quite small differences in both cases. In addition, the group mean MASK gestural profiles for the children were strikingly similar to the MASK and EMA profiles of the adult.

### 4.1 Identification of gestural landmarks in speech

For self-paced productions of /ida/ and /ila/, analyses focused on the identification of the gestural landmarks for the /d/ and /l/ consonants and the measures of distance, velocity and duration in order to investigate the kinematic profiles of /d/ and /l/. While /l/ is a kinematically more complex and later developing sound than /d/, we expected that the 8-12 year old children in the present study would have largely mastered both of these sounds. The anterior constriction of the tongue tip of /l/ is mastered early in speech development, a reason why most of the articulatory errors of /l/ that children produce, are omissions of the tongue body retraction gesture (/j/ for /l/) (Studdert-Kennedy & Goldstein, 2003). Moreover, looking at the physiological development of the vocal tract, it has been reported that the tongue growth curve is stabilized around the age of 5;6 (Vorperian et al., 2005).

Speech motor behaviors have traditionally been studied by examining spatial and temporal measures such as maximum velocities, maximum amplitudes and maximum durations associated with single articulatory movements (Smith et al., 2000, Van Lieshout & Moussa, 2000). Assessment of the kinematic parameters of the consonants /d, l/ and their interactions, revealed that the two consonants are characterised by similar kinematic profiles. This similarity can be explained because both are voiced sounds, one being a stop and the other a liquid, which has also a central constriction but lateral openings. The smaller movement range for /l/ compared to /d/ might be because the /d/ is an oral stop, requiring a full tongue tip occlusion at the alveolar ridge but this position may not be reached for /l/ in central location. The target closing movements of /l/ are slightly shorter in duration but with lower maximum velocity (because of the smaller amplitude), which perhaps reflects the more complicated motion pattern (central constriction with lateral margin drops of the tongue).

### 4.2 MASK in children

Mature speech production requires precise neuromotor control of some 100 muscles associated with the vocal tract, control is finally achieved through a prolonged developmental learning process that extends at least until late adolescence. Previous articulographic work has shown that children’s speech is controlled in larger, less specified units than adult speech; that adult rates of production are not reached until 16 years of age; that some sounds are acquired later in development (e.g. /r/’s and /l/’s); and that children’s speech productions are less consistent/more variable than adult productions (Smith, 2006). To date, the assessment of speech motor control has been limited in paediatric populations. Data is particularly lacking for tongue kinematics because of the experimental difficulties in obtaining non line-of-sight measurements within the oral cavity of children (Goozee et al., 2000; Cheng, Murdoch, & Goozée, 2007). For functional brain imaging, children pose additional difficulties due to the need to minimise head and body movements during the scans (Johnson et al., 2010).

The present study measured articulographic data concurrently with neuromagnetic brain activity in ten 8 - 12 year old children. With some prior training in a mock scanner (Rapaport et al, 2019) all children in this age group followed instructions well with minimal errors or interruptions: As a group they showed near-adult levels of compliance (see also Van Lieshout et al., 2002).

Most researchers have investigated variability of speech motor control by measuring the kinematic parameters of specific targeted consonants and they have utilised stimuli produced in a self-paced manner (Murdoch et al., 2012, Loh et al., 2005). Loh et al. (2005) used EMA to examine the variability of jaw movements in two children after traumatic brain injury (TBI) and in a control group of typically developing peers. Their experimental task consisted of reading aloud 24 sentences in a comfortable rate and loudness. On the same line, Murdoch et al. (2012) used EMA and focused on the developmental variability of the lip and the tongue during opening and closing movements in three groups of children aged 6-7, 8-11, 12-17 and adults. Their task consisted of two set of utterances which participants had to produce at their conversational rate and loudness. A part of the focus in Goffman & Smith’s study (1999) was to examine the stability of single open-close movements in children 4, 7 years old and adults using an Optotrak camera. The stimuli consisted of five minimal pairs which were embedded in a carrier phrase and participants were instructed to produce them with approximately 1s pause between each. Similarly, Smith & Goffman, (1998) used an Optotrak camera to measure the stability of speech movements in children and adults. They measured the amplitude and the peak velocity of two opening and two closing lip movements for *bob* and *pup* embedded in the phrase ‘Buy Bobby a puppy’ which participants had to produce in a normal rate and loudness.

We used the reiterated DDK /pataka/ to measure the spatiotemporal variability of the tongue tip, tongue body and tongue dorsum across repetitions. With one exception we found that the group of children had higher STI scores than the adult participant for all tongue sensors, reflecting a lower level of stability and consistency of the articulatory movements. These results are in agreement with previous studies using the STI to measure the variability of a single articulator from childhood to adulthood. Murdoch et al. (2012) who examined the stability of tongue-tip, tongue-body and lower lip over sentence repetitions to capture the articulation of targeted consonants, found that children’s sentence productions were characterized by more variability compared to adults and this variability reduced with age. Moreover, Goffman & Smith, (1999) investigated the variability of the lower lip from childhood to adulthood while participants produced a range of minimal pairs. The results showed that children had higher STI values compared to the adult and that these articulatory movements became systematically more stable with age.

### 4.3 Scientific and clinical implications

The development of the brain mechanisms of speech motor control remains an under-studied topic. In a review of this topic Smith (2010) stated: “One of the most astonishing conclusions one reaches after completing a review of this literature is that there is a real paucity of studies of oral-motor development for speech. There are very few laboratories doing work in this area…”. The author concludes that “…advances in understanding the neural control of speech motor development will depend on assembling … research teams who can make multi-levelled observation of both peripheral speech motor output and the neural activity generating that output, in children …. and adults”. MASK provides the capability for just such multi-levelled observations, and uniquely, within the same experimental setup.

**Figure 7** illustrates the capability of MEG neuroimaging for resolving spatial patterns of activity in the motor control regions of the brain corresponding to the known somato(motor)-topic representation of the human body, while **Figure 8** shows that distinct temporal-spectral profiles of brain activity are observed in different regions of the speech motor network. The robust beta and gamma oscillations of **Figure 8** are established neurophysiogical markers of anticipatory motor planning, response planning and selection of sensory targets in the adult brain (Cheyne, 2013). In our work on movement-related brain activity during manual motor tasks in children 3 to 5 years of age we have demonstrated that children show similar movement-evoked fields to those observed in adults, but significantly delayed (Cheyne et al., 2014). We also observed sensorimotor cortex mu (8–12 Hz) and beta (15–30 Hz) oscillations, but with different timing and mean frequency compared to adults. For example, high-frequency (70–80 Hz) gamma bursts were detected in the motor cortex at movement onset and correspond to that observed in adults. In a recent follow-up study (Johnson et al., 2019 preprint) we re-scanned a subset of these children two years after the original study. We found that the high gamma activity increased significantly in amplitude in the second scan; there were also significant changes in other neurophysiological markers including mu, beta, low gamma, and MFs).

**Figure 7.**
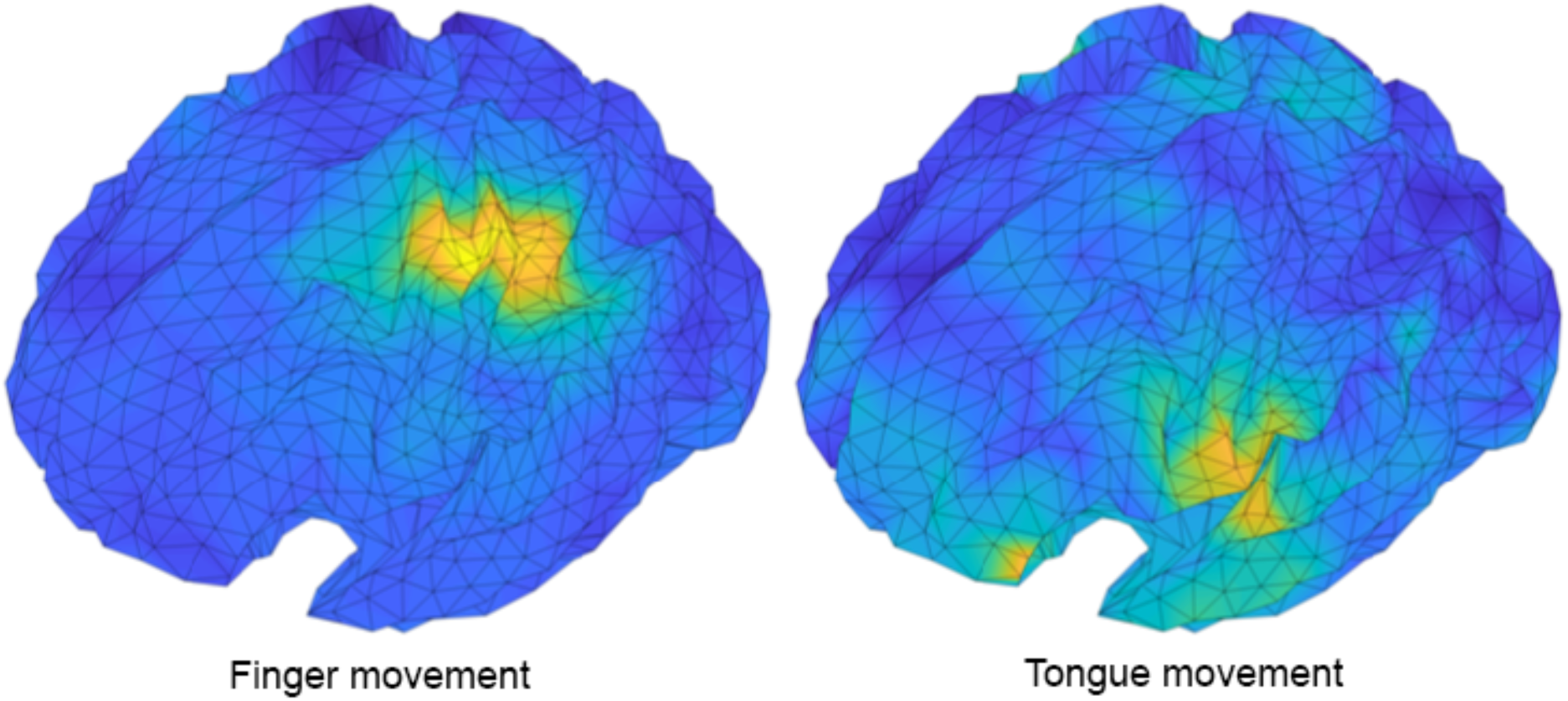
Group mean source locations for MEG brain activity recorded during self-paced right-hand finger taps (16 adult participants) and tongue tip movements during self-paced productions of /pataka/ (13 adult participants). Finger movements show activation in the dorsal hand sensorimotor region (Brodmann’s area 3 and 4) while tongue tip movements during /pataka/ productions are associated with activation of ventral sensorimotor speech cortex.

**Figure 8.**
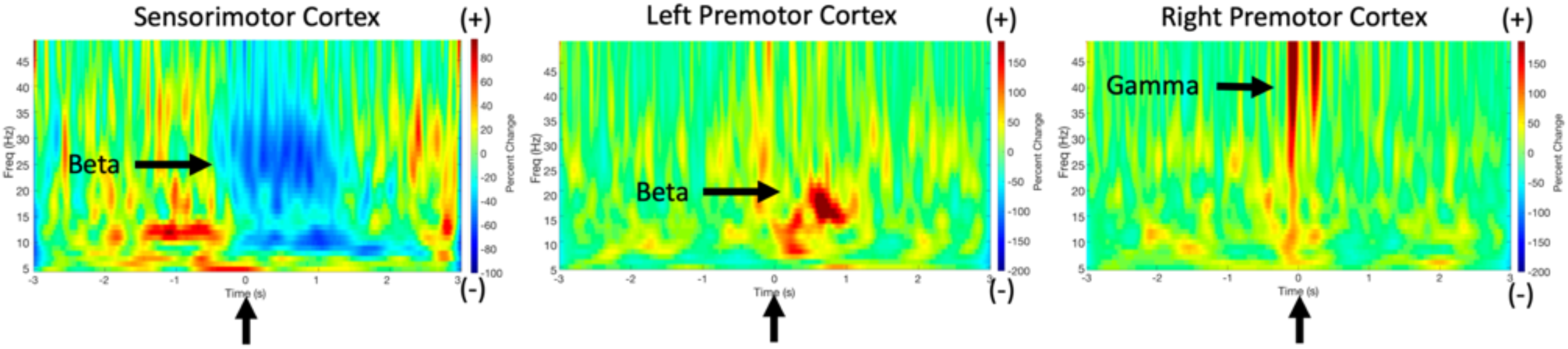
MEG responses to lip movements during productions of /pataka/. Source activity in sensorimotor speech cortex (Brodmann areas 3,4) in a single participant shows decreased beta-band activity prior to and during speech lip movements. Left premotor (Brodmann areas 44/45) source activity shows increased beta-band activity. Right premotor source shows transient bursts of gamma-band activity. Arrows indicate lip movement onset. Beamformer-based “virtual electrode” source profiles are highly robust to speech movement and other artefactual sources (Cheyne et al., 2007).

The problem of how humans develop speech is a central, unanswered question of neurolinguistics and movement science. The topic is manifestly under-studied (Smith, 2010). Answers to this problem will inform and constrain theoretical models of language, and will have practical implications for our understanding and treatment of speech-language pathologies. These include the now well-replicated finding of a greater incidence of comorbid motor coordination and planning problems in children with language impairments (Iverson and Braddock, 2011). The development of speech motor control is also highly relevant to the burgeoning fields of speech prosthetics and brain computer interfaces. Recent invasive electrocorticographic (ECoG) studies of human patients have demonstrated that the kinematic parameters measured by our MASK setup are precisely those that are encoded and mapped in speech motor cortex (Chartier et al., 2018). Further, a recent landmark ECoG study demonstrates that neurophysiological measurements of these parameters can be decoded directly into intelligible speech (Anumanchipalli et al., 2019).

### 4.4 Future developments

Since the collection of the present data we have incorporated electronics for real-time processing and display of both the speech tracking positions and the MEG brain data. As also used in a typical EMA setup, the real-time display greatly improves overall tracking data quality by providing real-time feedback on the patency of each tracking sensor and allowing investigators to correct or replace problematic sensor attachments during the recording session.

### 4.5 Conclusions

The present results demonstrate that the MASK technique can be used to reliably characterise movement profiles and kinematic parameters that reflect development of speech motor control, while simultaneously measuring the brain activities that provide this control. MASK brings articulography into the brain itself and thereby bridges a crucial methodological gap between the fields of speech science and cognitive neuroscience. The importance of this gap has recently been emphasised by invasive ECoG studies which have demonstrated that speech motor cortex operates by encoding and computing speech kinematic parameters that can be derived only with detailed measurements of the movements of individual articulators, including non line-of-sight measurements of the oral cavity. This new capability sets the stage for new cross-disciplinary efforts to understand the neuromotor control of human speech production.

## Acknowledgements

This research was supported by Australian Research Council Discovery Project Grant DP170102407 to Blake Johnson, Douglas Cheyne and Pascal van Lieshout and Natural Sciences and Engineering Research Council CHRP Grant # 385824-10 to Douglas Cheyne, Pascal van Lieshout and Tom Chau.

## Author Contributions

Blake Johnson, Qingqing Meng and Ioanna Anastasopoulou contributed equally to this work. Blake Johnson, Douglas Cheyne, Pascal van Lieshout, Qingqing Meng, Ioanna Anastasopoulou and Michael Proctor conceived and designed the study. Cecelia Jobst contributed to the analysis and interpretation of the MASK and MEG data. Louise Ratko and Tunde Szalay contributed to acquisition, and analysis of the EMA data. All authors contributed to the drafting of the work and revising for critically important intellectual content, provided final approval of the version to be published, agreed to be named on the author list and approved of the full author list.

